# The nutritional environment is sufficient to select coexisting biofilm and quorum-sensing mutants of *Pseudomonas aeruginosa*

**DOI:** 10.1101/2021.09.01.458652

**Authors:** Michelle R. Scribner, Amelia C. Stephens, Justin L. Huong, Anthony R. Richardson, Vaughn S. Cooper

## Abstract

The evolution of bacterial populations during infections can be influenced by various factors including available nutrients, the immune system, and competing microbes, rendering it difficult to identify the specific forces that select on evolved traits. The genomes of *Pseudomonas aeruginosa* isolated from the airway of patients with cystic fibrosis (CF), for example, have revealed commonly mutated genes, but which phenotypes led to their prevalence is often uncertain. Here, we focus on effects of nutritional components of the CF airway on genetic adaptations by *P. aeruginosa* grown in either well-mixed (planktonic) or biofilm-associated conditions. After only 80 generations of experimental evolution in a simple medium with glucose, lactate, and amino acids, all planktonic populations diversified into lineages with mutated genes common to CF infections: *morA*, encoding a regulator of biofilm formation, or *lasR*, encoding a quorum sensing regulator that modulates the expression of virulence factors. Although mutated quorum sensing is often thought to be selected *in vivo* due to altered virulence phenotypes or social cheating, isolates with *lasR* mutations demonstrated increased fitness when grown alone and outcompeted the ancestral PA14 strain. Nonsynonymous SNPs in *morA* increased fitness in a nutrient concentration-dependent manner during planktonic growth and surprisingly also increased biofilm production. Populations propagated in biofilm conditions also acquired mutations in loci associated with chronic infections, including *lasR* and cyclic-di-GMP regulators *roeA* and *wspF*. These findings demonstrate that nutrient conditions and biofilm selection are alone sufficient to select mutants with problematic clinical phenotypes including increased biofilm and altered quorum sensing.

**Importance:** *Pseudomonas aeruginosa* produces dangerous chronic infections that are known for their rapid diversification and recalcitrance to treatment. We performed evolution experiments to identify adaptations selected by two specific aspects of the CF respiratory environment: nutrient levels and surface attachment. Propagation of *P. aeruginosa* in nutrients present within the CF airway was alone sufficient to drive diversification into subpopulations with identical mutations in regulators of biofilm and quorum sensing to those arising during infection. Thus, the adaptation of opportunistic pathogens to nutrients found in the host may select mutants with phenotypes that complicate treatment and clearance of infection.

## Introduction

During infection, a subset of spontaneous mutations in the pathogen population may produce dangerous clinical consequences including resistance to antimicrobial treatment and immune clearance (1–4). These become more probable with increasing generations of growth as populations expand or persist (5–7). It is therefore critical to characterize adaptations that are selected *in vivo*, the causative mutations, and the environmental factors that contribute to their selection. Longitudinal *Pseudomonas aeruginosa* isolates from many patients with cystic fibrosis (CF) share common evolved traits, including high biofilm phenotypes (8), increased antimicrobial resistance (9), loss of O-antigen (10), and increased alginate production (11). These traits also often vary among isolates within patients and this diversity is thought to contribute to ineffective clearing by treatment (12, 13) and ultimately increased morbidity and mortality (14, 15). Whole-genome sequencing (WGS) of these isolates has identified convergent mutations in certain genes that persist within the CF respiratory environment (2, 8, 16). One of the most commonly mutated genes is *lasR*, encoding a transcriptional regulator of quorum sensing, and mutations in regulators of biofilm production are also common, among others (2, 17, 18). However, numerous factors could select for phenotypic and genotypic diversity in the CF airway (12, 19), making it challenging to infer causes of the prevalence of these mutations and phenotypes.

Evolution experiments in models of host systems can clarify which factors most influence pathogen fitness *in vivo*. The nutritional environment is increasingly appreciated as a major selective pressure, which motivated development of an artificial sputum medium that approximates concentrations found in patients with CF and produces similar growth phenotypes and transcriptional responses as actual sputum (20–22). Evolution experiments with *P. aeruginosa* have been conducted in this medium to identify beneficial mutations in particular conditions, including antibiotic pressure (23), biofilm lifestyle (24), and presence of mucins (23, 25). Several observations are consistent across these studies, particularly the rapid diversification of biofilm, motility, and colony phenotypes. Mutations in certain genes have also been identified repeatedly including *lasR, wsp*, and flagella synthesis genes (23, 24, 26, 27).

Because these common mutations produce broad pleiotropic effects, determining which phenotypes were initially adaptive within the host remains unresolved. For instance, the *lasR* gene encodes the transcriptional regulator of the LasRI acyl-homoserine lactone (AHL) quorum sensing system which regulates the expression of hundreds of genes, including other quorum sensing systems (RhlRI and PQS). Loss-of-function *lasR* mutations reduce production of extracellular proteases (28), improve fitness in microoxia (29), increase resistance to ceftazidime (24), improve growth in amino acids (30), and produce a “lysis and sheen” colony morphology (30). LasR mutants are frequently described as social cheaters because they can use public goods such as proteases produced by competitors without undergoing the cost of producing these products themselves (31, 32), although recent findings suggest that *lasR* mutants actually overproduce these public goods when co-cultured with a LasR+ strain (33). In addition, mutations in regulators of cyclic-di-GMP, a second messenger that promotes biofilm and suppresses motility when upregulated, may also produce a wide range of effects on cell cycle, virulence, and motility (34, 35). These mutations may be selected for the complete suite of new phenotypes they cause or perhaps only one of them. Modeling specific subsets of the infection environment is therefore required to quantify contributions of different selective pressures to pathogen evolution.

We sought to identify which adaptations to the CF airway may be explained exclusively by specific nutrients by performing an evolution experiment of *P. aeruginosa* strain PA14 in a defined medium containing carbon sources prevalent in this environment. Replicate populations were propagated either by serial broth dilution or in a biofilm model using beads to study the influence of lifestyle on mutant selection (36). We performed whole population genome sequencing of each lineage and sequenced isolated clones to infer population structure and to link evolved phenotypes to genotypes. We also characterized the consequences of these mutations on clinically relevant phenotypes including biofilm production and motility to examine how these adaptations could influence pathogenesis.

## Results

To examine the role of nutrients in *P. aeruginosa* adaptation to the CF airway, strain PA14 was propagated in substrates found to be prevalent in the cystic fibrosis respiratory tract (11mM glucose, 10mM LD-lactate, and 4mM amino acids) for 12 days. The concentrations of these carbon sources differ from those in an established synthetic CF medium (SCFM; 3.2mM glucose, 9.3mM lactate, and 19mM amino acids) (21), but were used to facilitate comparisons with previous studies by our group and are within the range previously reported for CF sputum with the exception of increased glucose (21, 37). Populations were propagated with either planktonic selection through a 1:100 dilution into fresh medium every 24 hours or biofilm selection by transferring a colonized polystyrene bead to fresh medium every 24 hours as previously described (38). Although other factors associated with our culture conditions also influence selection, we can compare adaptations in this medium with those using different medium but identical transfer protocols to identify mutations associated with growth in CF nutrients (39). Population transfer sizes were ~10^8^ cells in both planktonic and biofilm transfer methods, producing approximately 6.7 generations/day and 80 generations over the course of the experiment. We estimate that approximately 10^6^ new mutations occur each day in this system based on reported mutation rates of this and related strains (39–42), but these large effective population sizes ensure that only the fittest mutants within each population will rise to a detectable frequency within this short time frame (37). Thus, the mutations reported here almost certainly produce adaptations.

### Genetic targets of selection in CF nutrients mimic *in vivo* adaptation

We performed whole genome sequencing of five evolved planktonic and five evolved biofilm populations to read depths of 86-254X to identify selected genotypes (Figure 1). After only twelve days, mutations in *lasR* (PA14_45960) and *morA* (PA14_60870) evolved within nearly every planktonic population. Mutations in *lasR* included large deletions encompassing both *lasR* and *lasI* genes (PA14_45920 to PA14_45440, and PA14_45800 to PA14_46240), nonsynonymous SNPs, and nonsense mutations (Figure 2A). The *lasR* and *lasI* genes encode the regulator and autoinducer synthesis proteins, respectively, of the Las quorum sensing system (43). The *morA* gene encodes a protein with both diguanylate cyclase (DGC) and phosphodiesterase (PDE) domains, which produce and degrade second messenger cyclic di-GMP, respectively (35). Nonsynonymous substitutions occurred at five different residues within *morA* (A1109V, K1123E, N1124K, E1153K, and L1155Q), indicating that they arose independently and were selected in parallel (Figure 2B). Mutations in *morA* and *lasR* are two of the most frequently mutated genes in CF clinical isolates (2, 8, 30), and strikingly, this experiment selected mutations in some of the same residues altered during CF infections, including *lasR* P117 and A231 (8). We also detected multiple instances of residue and domain-level parallelism between *morA* mutations from this experiment, from other evolution experiments in SCFM or rich media, and from clinical isolates (Figure 2B, Figure S1) (2, 23, 44). This convergence demonstrates that the nutrients found in CF sputum are sufficient to select for the same mutations identified in clinical isolates.

**Figure 1.**
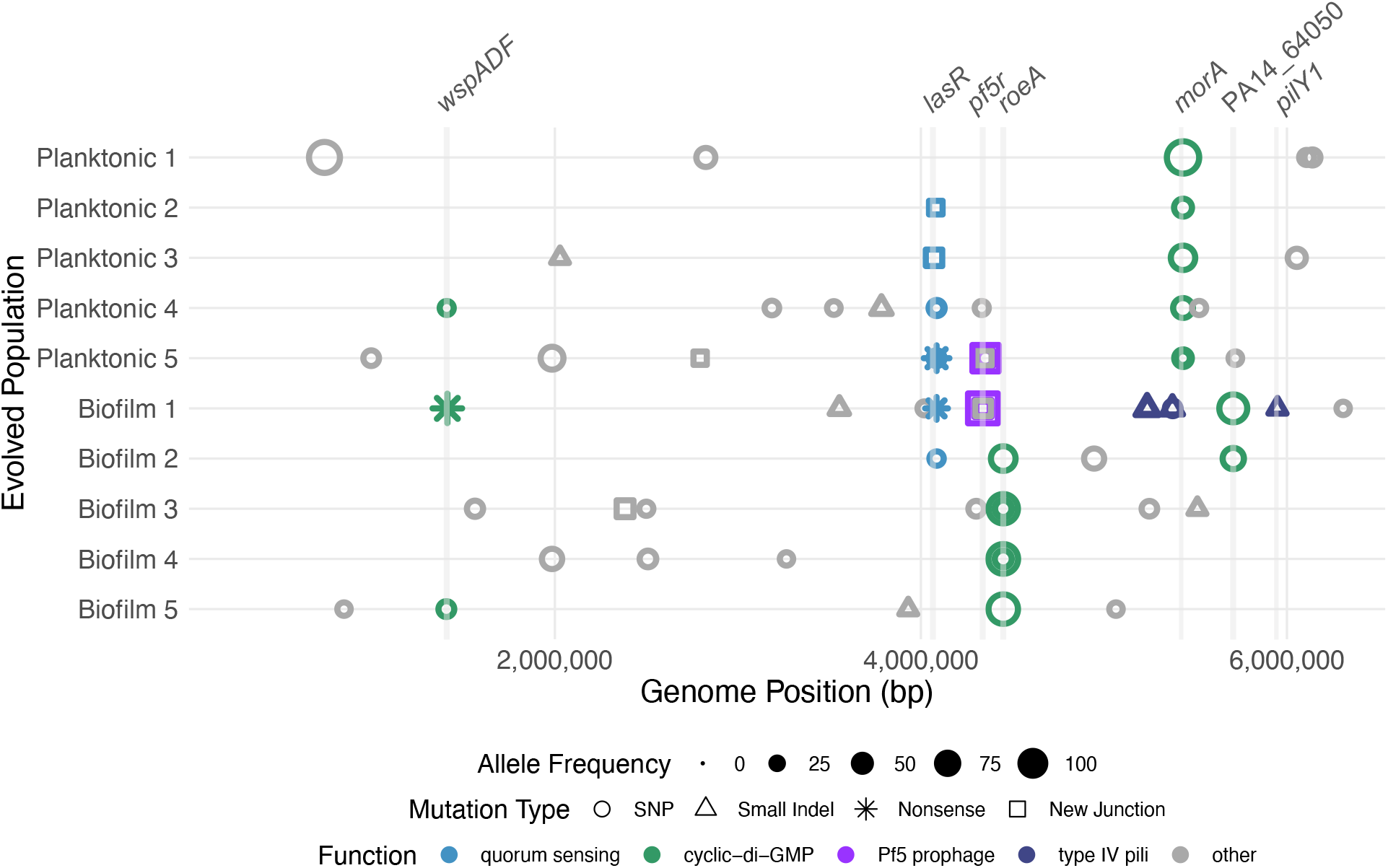
Propagation of *P. aeruginosa* in nutrients present in the CF airway rapidly selects for mutations in regulators of quorum sensing and cyclic-di-GMP. Mutations were inferred from whole genome sequencing of evolved populations after 12 days of selection. All mutations detected by population WGS are indicated and genes that acquired multiple mutations are labeled. Details of detected variants are in Supplementary Data Table 1. Allele frequency for each mutation is indicated by symbol size, mutation type by symbol shape, and function of the impacted loci by symbol color. Mutations detected by new junction evidence, which include insertions, large deletions, and structural rearrangements, are indicated at the position of the most upstream side of the junction.

**Figure 2.**
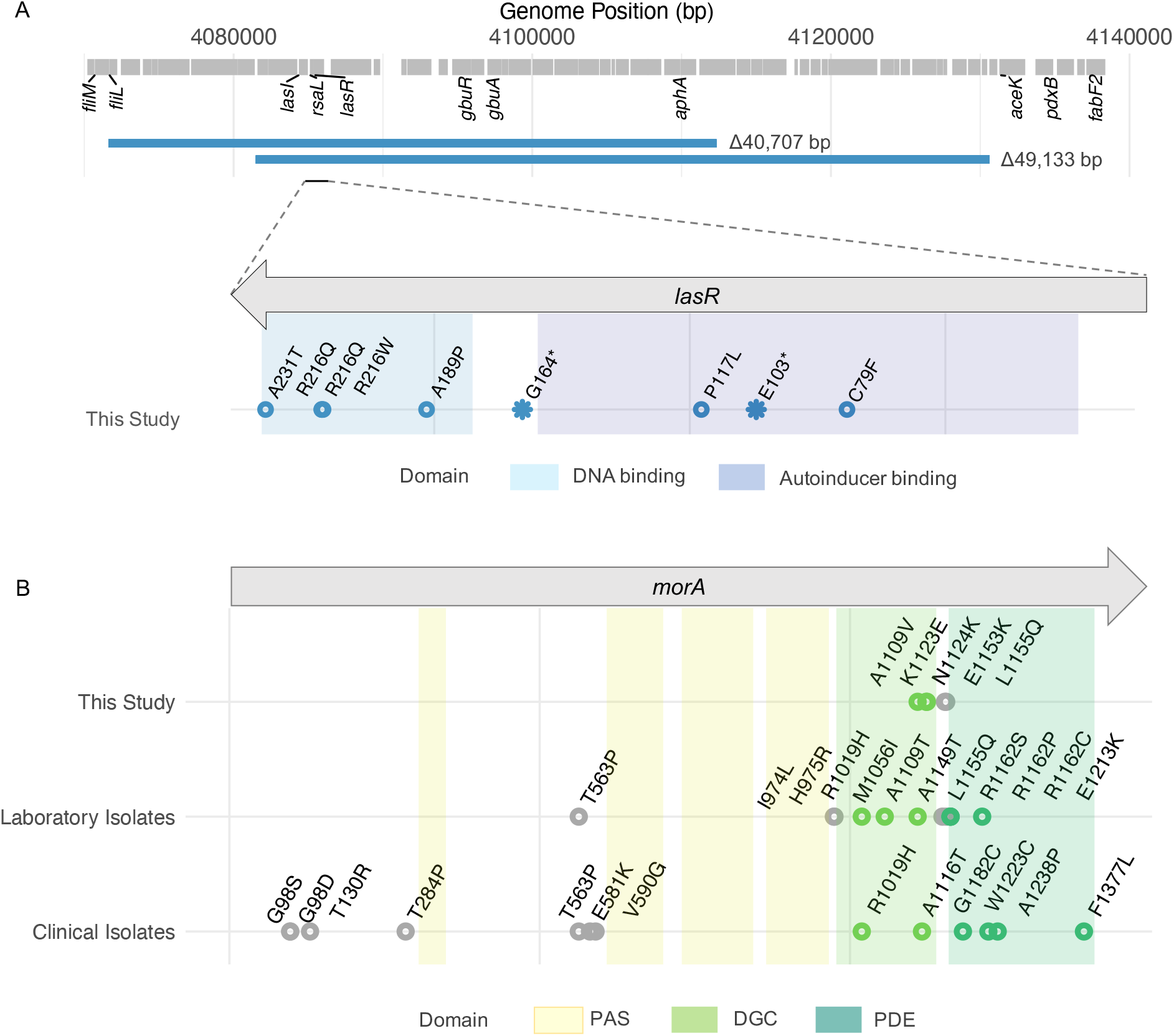
Convergent evolution of mutations in *lasR* and *morA* reveal sites under strong selection. A) Two large deletions that include the *lasI* and *lasR* genes were detected as well as many nonsynonymous SNPs and missense mutations. Top: the extent and overlapping region of these deletions; below: evolved *lasR* mutations by relative position. B) Mutations in *morA* were also frequently selected, primarily within the diguanylate cyclase (DGC) domain and linker region between the DGC and phosphodiesterase (PDE) domains (This Study). We also identified *morA* mutations in other evolution experiments in our laboratory and other studies (Laboratory Isolates) (23, 44). Identical mutated sites have been identified in clinical isolates from patients with cystic fibrosis (Clinical Isolates) (2, 8).

Biofilm-adapted populations also nearly exclusively selected mutations associated with cystic fibrosis infection including genes regulating cyclic-di-GMP, quorum sensing, and type IV pili (2, 4). All biofilm populations acquired at least two mutations in regulators of cyclic-di-GMP (*wspAF, roeA*, and PA14_64050), often on different lineages at high frequencies (Figure 1). These findings add to evidence that modulating levels of cyclic-di-GMP is advantageous in a biofilm environment (45, 46). Yet, the PA14 genome harbors ~40 genes containing DGC domains, PDE domains, or both that could alter cyclic-di-GMP levels (47), so the repeated selection of mutations in only these three genes suggests that these regulators of cyclic-di-GMP are not redundant but rather are environment-specific, as shown elsewhere (48). Biofilm lineages 1 and 2 also acquired mutations in *lasR*, suggesting that altered quorum sensing was beneficial in both planktonic and biofilm environments. However, identical evolution experiments using different growth medium containing arginine as sole carbon source did not select for mutations in *lasR* or these cyclic-di-GMP regulators, indicating that their benefit is linked to these nutrients (39). Most importantly, the almost exclusive selection of mutations known to arise during chronic infection in CF suggests that our simple nutritional model closely recapitulates several major selective forces acting in the CF airway.

### Mutations in *lasR*, *morA*, and *wspF* underlie rapid morphological and phenotypic diversification

Colony morphologies became conspicuously diverse in both biofilm and planktonic treatments after twelve days of selection, so we picked clones with representative phenotypes and sequenced their genomes to identify the causative mutations (Figure S2). These genotypes demonstrated that *lasR*, *morA*, and *wspF* lineages arose independently and identified additional *lasR* and *wspF* mutations that were undetected by population WGS. Mutations in *wspF* or between the DGC and PDE domains of *morA* produced wrinkly or rugose small colony variants (RSCVs). We measured the motility and biofilm production of isolated mutants and found that *lasR* and *morA* mutants from the same population were functionally distinct (Figure 3). All *morA* mutations decreased swimming motility and increased biofilm, whereas *lasR* mutants showed no change in biofilm or swimming motility, except for one that also had disrupted flagellar genes (*fliM* and *fliL)* which lost swimming motility. Unsurprisingly, all mutants isolated from biofilm-evolved populations produced more biofilm and most showed decreased swimming motility (Figure 3) and swarming motility (Figure S3). Therefore, similar phenotypes evolved in replicate populations associated with parallel genotypes.

**Figure 3.**
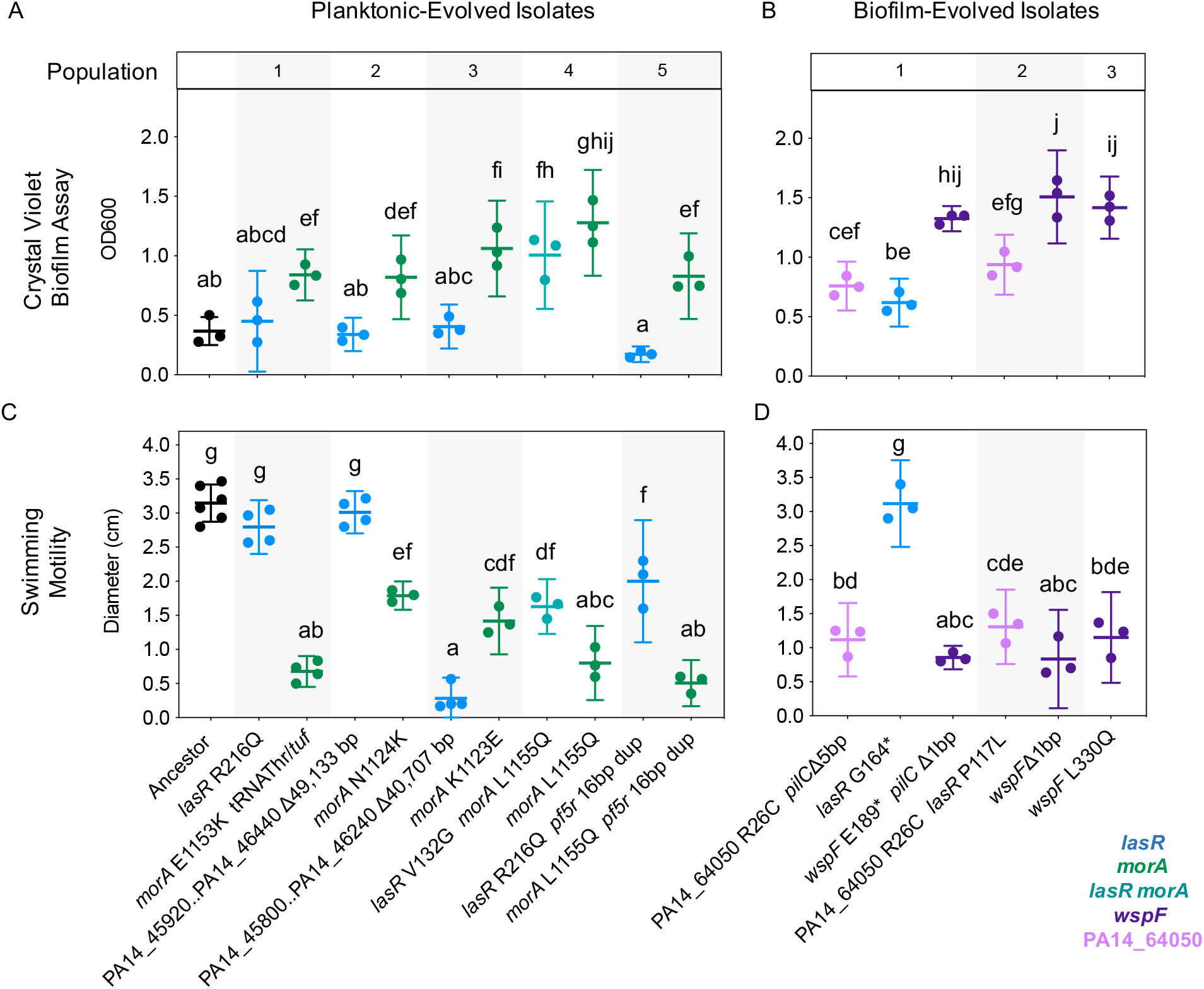
Mutations in *lasR* and in multiple cyclic-di-GMP regulators produce coexisting subpopulations with distinct biofilm and motility phenotypes. Biofilm production by isolates from A) planktonic lineages and B) biofilm lineages. Swimming motility by isolates from C) planktonic lineages and D) biofilm lineages. Strains are labeled by genotype. Data points represent the average of technical replicates from at least three independent experiments and are shown with mean and 95% CI. Data were analyzed by one-way ANOVA (Biofilm: F(16,34) = 32.76, p < 0.0001, Swimming Motility: F(16,41) = 72.67, p < 0.0001) with Tukey’s multiple comparisons test. Groups labeled with the same letter are not statistically different (p < 0.05).

### *morA* mutations produce nutrient concentration-dependent increases in fitness

The repeated selection of *morA* and *lasR* mutations in planktonic populations indicates that independently of any advantages of increased biofilm or altered virulence factor production in a host, they produce growth advantages. Prior studies have shown that *morA* mutant phenotypes depend on carbon sources (48) and that *lasR* mutants enhance growth in amino acids (30). To build upon these findings and to explore possible mechanisms explaining their coexistence, we measured mutant fitness in each of the carbon sources in the evolution medium: glucose, amino acids, and lactate (Figure 4). A mutant with a nonsynonymous SNP in the linker domain of *morA* outcompeted the ancestor in medium containing all carbon sources but was less fit in media containing only a subset of the carbon sources (Figure 4B). Other Isolates with SNPs in *morA* also grew worse than the ancestor when cultured in only amino acids or only lactate (Figure S4). These findings suggest that fitness of *morA* mutants may be influenced by nutrient identity, nutrient concentration, or both. To distinguish between these possibilities, we performed competition assays in various nutrient levels and found that *morA* mutant fitness was indeed dependent on nutrient concentration (Figure 4B). In fact, halving the concentration of each of the carbon sources in the evolution medium eliminated the growth advantage of *morA* over WT. Therefore, nutrient abundance alone is sufficient for the selection of *morA* mutants, though nutrient composition may modulate the strength of this selection.

**Figure 4.**
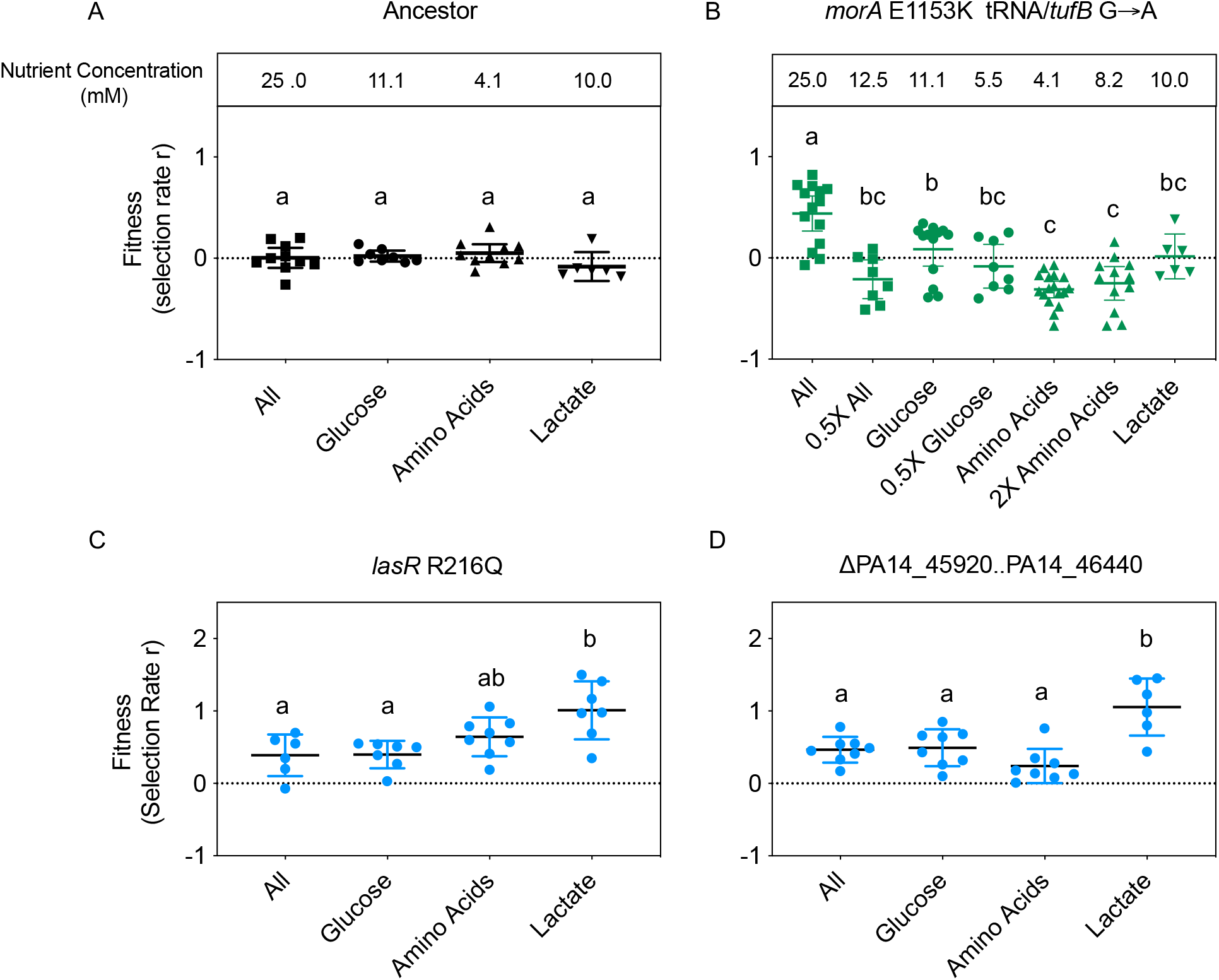
Environment-specific fitness advantages of *lasR* and *morA* mutants. A marked ancestral strain was competed against A) the ancestral strain, B) an isolate with a *morA* SNP (E1153K) and an intergenic SNP between a tRNA and the *tufB* gene, C) an isolate with a SNP in *lasR* (R216Q), and D) an isolate with a large deletion encompassing *lasR* and *lasI*. Competitions were performed in the medium used for the evolution experiment (All) as well as medium containing only a subset of the carbon sources (Glucose, Amino Acids, Lactate). The *morA* mutant was also competed in media in which the concentration of nutrients was doubled or halved to determine the effect of nutrient concentration on fitness (0.5X All, 0.5X Glucose, 2X Amino Acids). Data was collected from at least two independent experiments. Mean and 95% CI are shown. Within each genotype, statistical differences in fitness in each media were determined using ANOVA (Ancestor: F(3,30)=1.587, p=0.2131, *morA* tRNA/*tufB*: F(6,71)=14.96, p<0.0001, *lasR* R216Q: F(3,26)=11.14, p<0.0001, ΔPA14_45920..PA14_46440: F(3,24)=6.608, p=0.0021) with Tukey’s multiple comparisons test. Groups labeled with the same letter are not statistically different.

### *lasR* mutations are beneficial in the absence of a competitor

The fitness advantage of *lasR* mutants in chronic infections has been proposed to derive from multiple features, including social cheating, growth advantages in amino acids or in microoxic environments, and increased growth at high cell densities (28–31, 49–52). Social cheating refers to the ability of a *lasR* mutant to reap the benefits of public goods secreted by LasR+ cells within the population, including proteases, pyocyanin, and hydrogen cyanide, without undergoing the cost of producing those factors themselves. However, we observed that in the absence of a competitor, *lasR* mutants demonstrated greater area under growth curves than the ancestor (AUC, Figure S4) and similar or greater growth yields (Figure S5). This result shows that exploitation of a competitor is unnecessary for increased *lasR* mutant fitness.

Prior studies of *lasR* mutants have implicated altered carbon catabolite repression as the source of their growth advantages in aromatic amino acids (30). Yet we found that *lasR* mutants attained similar fitness advantages in media containing only glucose or only lactate as in only amino acids (Figure 4C and 4D). Nonetheless, we tested whether *lasR* mutations altered carbon catabolite repression by analyzing the nutrient levels in the spent medium of a *lasR* mutant compared to spent medium of the ancestor. We cultured strains separately in the evolution medium and used HPLC to quantify nutrient levels every hour for the first eight hours of growth. We hypothesized that *lasR* mutations may alter the rate or order of nutrient consumption when growing in medium containing multiple carbon sources. We noticed *lasR* mutants produced a spike in amino acid levels at one hour post inoculation that we cannot explain, but no subsequent significant differences in nutrient consumption rate or order were found (Figure S6). Therefore, we cannot attribute the selective advantage of *lasR* mutants in this complex medium to altered carbon catabolite repression, but it is possible that some relevant metabolic changes were too subtle to be detected using this approach. The specific metabolic source of the growth advantage of *lasR* mutations therefore remains unclear in this environment and merits further study.

### Ecological interactions between *morA* and *lasR* mutants facilitate the maintenance of diversity

Both *morA* and *lasR* genotypes became prevalent in every planktonic population and defined coexisting subpopulations in four out of five lineages, prompting us to ask if their relative fitness advantages derived from ecological interactions between them. We tested their ability to facilitate each other’s coexistence by mixing genotypes at different starting concentrations (Figure 5). Each mutant was more fit when introduced at a lower proportion, consistent with a negative frequency-dependent interaction, or a relationship in which the fitness of a competitor increases as it becomes rarer (53, 54). We repeated this experiment using different *morA* and *lasR* mutants and found this frequency-dependent interaction was consistent (Figure S7). Therefore, nutrients present in the cystic fibrosis lung environment are sufficient to rapidly drive stable, functional diversification, even in the absence of spatial heterogeneity. This finding is highly relevant to clinical settings as diversity within infections may increase recalcitrance to treatment (13).

**Figure 5.**
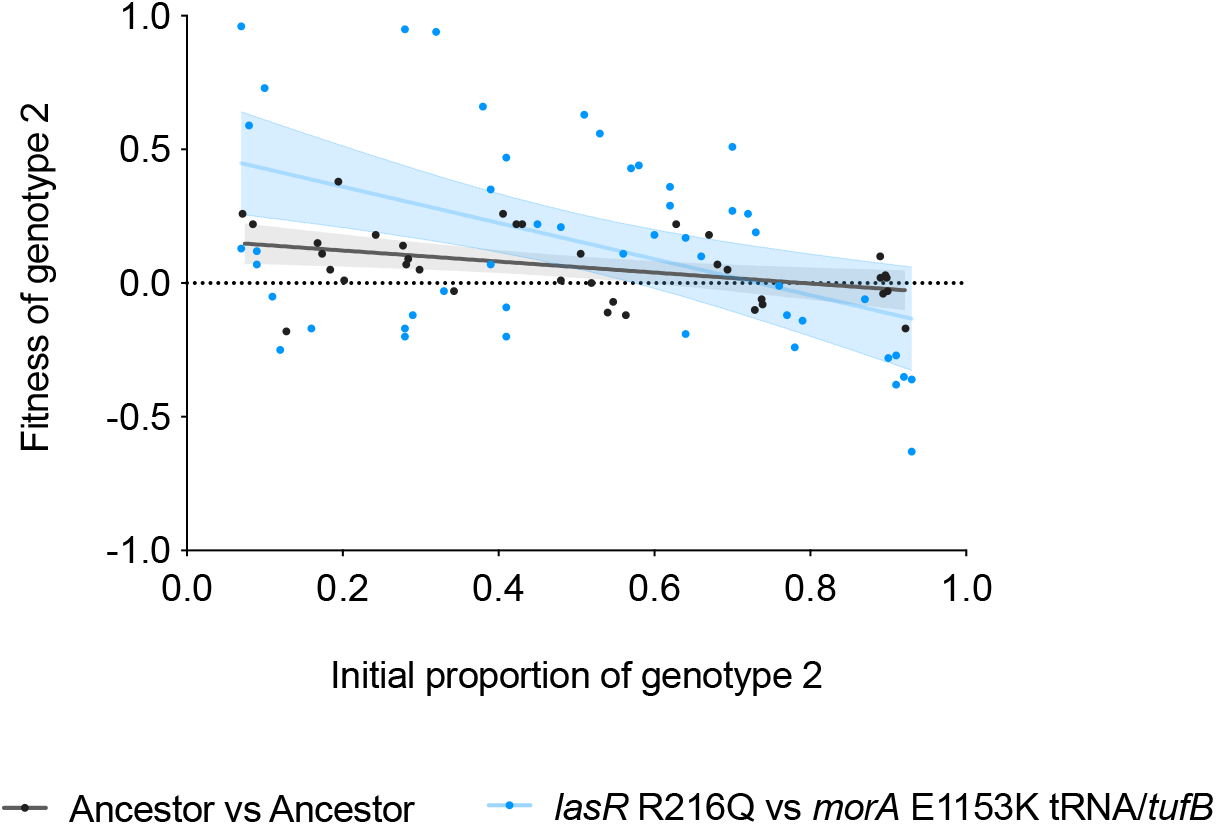
Relative fitness of *lasR* and *morA* mutants is greater when in the minority. We performed direct competitions at the starting concentrations indicated and measured relative fitness at 48 hours. Genotypes were differentiated using a lac-marked ancestral strain for the ancestral competitions and by colony morphology for the *lasR* mutant versus *morA* mutant competitions. The first competitor listed is genotype 1, the second is genotype 2. Data was analyzed by linear regression, shaded area represents 95% CI of the regression. Ancestor versus ancestor: slope = −0.2060, *lasR* mutant versus *morA* mutant: slope = −0.6760, difference between the slopes was statistically significant F = 4.195, DFn = 1, DFd = 83, p=0.0437.

## Discussion

The realization that chronic infections of the CF airway, wounds, and other sites by *P. aeruginosa* not only involve adaptive evolution, but also a process of diversification, has motivated numerous explanations (19). Most of these understandably focus on aspects of ourselves as hosts. However, one of the most basic and essential components of host-microbe interactions is nutrition. Here we show that both adaptation and diversification of *P. aeruginosa* can be explained solely by a subset of nutrients found in the cystic fibrosis respiratory environment at high concentrations. Moreover, selected genotypes acquire mutations identical to those recovered from infections, including mutations in cyclic-di-GMP regulators (*morA, wspF, roeA)* and quorum sensing regulators (*lasR*). Further, the phenotypes caused by these mutations are of potential clinical importance. Loss-of-function *lasR* mutations have been shown to alter the production of virulence factors and increase resistance to beta-lactam antibiotics (24, 30), and are associated with lung disease progression (55). Similarly, high biofilm phenotypes such as those produced by *morA* mutants (RSCVs) have been shown to increase persistence under stresses like the immune system and antibiotics, making these infections more difficult to treat (26, 56–58). Selection for either of these phenotypes alone merits concern, but here we show that *morA* and *lasR* mutants frequently are co-selected *in vitro* and facilitate one another’s coexistence. Though the mechanism underlying this ecological relationship is yet to be identified, the maintenance of this diversity poses its own risk for population persistence in the face of treatment.

One defining phenotype of many *P. aeruginosa* infections is increased secreted polysaccharides such as those produced by *morA* mutant genotypes. Their selection during planktonic serial transfer was surprising because biofilm matrix components ought to be costly to produce. Yet our results suggest that although the selected *morA* mutations increase biofilm, they were likely selected due to their strong fitness advantage when nutrients are abundant. In recent work, Katharios et al. suggested a model by which the phosphodiesterase activity of *morA* is induced during nutrient limitation to repress costly biofilm production and thus prevent cell death (59). Consequently, disrupted *morA* signaling results in increased biofilm during nutrient abundance, but cell death during nutrient limitation. Our findings are consistent with this report: nonsynonymous substitutions in *morA* increased biofilm but were costly for fitness when nutrient levels were reduced. Together our results suggest that the nutrient abundance within the cystic fibrosis lung environment plays an important role in the selection of high biofilm mutants.

Mutants of *lasR* have been recovered even more frequently from infections and studied intensively. A common model is that these genotypes can use public goods (i.e. diffusible, non-sequestered products) produced by competitors without engaging in the costs of quorum sensing itself (31, 60, 61). However, we find that these mutants achieve equivalent or higher net growth than the ancestor when growing alone, demonstrating that social cheating is not required for their fitness advantage in these conditions. Metabolic shifts conferred by *lasR* mutants have also been shown to contribute to fitness advantages in CF sputum, particularly increased fitness in amino acids (29, 30, 52). We were unable to detect altered carbon catabolite repression for *lasR* mutants by HPLC of spent media or amino acid-dependent fitness advantages in our experimental conditions, but *lasR* mutations may have produced metabolic changes that we were unable to detect using these methods. Others have suggested that derepressed growth of *lasR* mutants at high cell densities may be responsible for their increased growth yields (30) and have shown that *lasR* mutants are resistant to cell lysis at high cell densities (51). This idea is compatible with our data as it would explain both increased growth yield in the absence of LasR+ competitors and increased fitness in the presence of competitors in media lacking amino acids. Advantage at high cell densities is also consistent with the frequent selection of *lasR* mutants in a broad range of environments including CF (2, 8), models of CF infection (24, 27), and laboratory media (27). However, the mechanisms underlying the fitness advantages conferred by these *lasR* mutants remain unclear.

This evolution experiment also reveals at least two curious findings regarding the genetics of adaptation by *P. aeruginosa*. First, adaptation to CF nutrients is produced by mutated global regulators of multicellular behavior (quorum sensing, biofilm formation) with remarkable consistency. Although many other ways of increasing fitness in these nutrients and propagation methods surely exist, it is notable that the most beneficial mutants in this environment were those which alter massive and complex regulons, rather than downstream effectors of cell behavior (34, 62). Second, we detected several large deletions and structural variants in prevalent genotypes, including those that result in the loss of both *lasI* and *lasR.* In clinical isolates, SNPs within *lasR* are frequently detected, however we hypothesize that large deletions may be underreported due to the computational challenge of identifying them in draft genomes produced by short-read sequencing. One-fourth of mutations in this evolution experiment were indels or structural variants, so sequencing efforts of clinical isolates ought to analyze these possible alterations whenever feasible.

In summary, extensive genetic adaptation and diversification is a well-known feature of *P. aeruginosa* populations growing in the cystic fibrosis respiratory tract (12) and experimental models of infection but the specific selective causes have been elusive (21, 24, 25, 63). Studying adaptation *in vivo* is challenging because obtaining an early infecting strain that serves as an appropriate internal reference for subsequent isolates is often not possible. Further, in more established infections, evolved mutations may be too numerous to infer the phenotypic impact any single change. Thus, linking mutations known to occur *in vivo* with the conditions that selected them and the phenotypes they confer is critical to understand how *P. aeruginosa* adapts to the CF respiratory environment. Remarkably, the evolution of genotypes producing more biofilm and with altered quorum sensing can be selected by a subset of nutrients found in the cystic fibrosis airway and coexist by reciprocal ecological facilitation. Although the presence of other microbes, host factors, or antibiotic pressure may only add to the potential diversification of opportunists like *P. aeruginosa*, they may not be required to promote increased persistence *in vivo*.

## Methods

### Evolution Experiment

*Pseudomonas aeruginosa* strain UCBPP-PA14 (64) was propagated for twelve days in minimal media supplemented with the nutrients shown to be abundant in the cystic fibrosis respiratory environment (21). The evolution experiment was described previously in Scribner et al 2020 (38), in which these lineages were used to test whether mutations in lineages exposed to tobramycin did not occur in the absence of antibiotic selection, but were not analyzed further. Briefly, the minimal medium consisted of 11.1mM glucose, 10mM DL-lactate (Sigma-Aldrich 72-17-3), 20 mL/L MEM essential amino acids, 10 mL/L MEM nonessential amino acids (Thermofisher 11130051, 11140050), an M9 salt base (0.1 mM CaCl_2_, 1.0 mM MgSO_4_, 42.2 mM Na_2_HPO_4_, 22 mM KH2PO_4_, 21.7mM NaCl, 18.7 mM NH_4_Cl), and 1 mL/L each of Trace Elements A, B, and C (Corning 99182CL, 99175CL, 99176CL). We propagated cultures in 18×150mm glass tubes containing 5mL of media and incubated in a roller drum at 150rpm at 37°C. Lineages were initiated by resuspending a single ancestral clone in PBS and using this suspension to inoculate each population. Five lineages each were propagated with either planktonic and biofilm selection: planktonic through a dilution of 50uL into 5mL of fresh media every 24 hours, and biofilm through transfer of a colonized bead to a tube with fresh media and 3 new beads. The evolution experiment continued for twelve days with sampling for freezing in 25% glycerol at −80 °C on days six and twelve. Planktonic populations were sampled by freezing 1mL aliquots and biofilm populations were sampled by sonicating a bead in PBS and freezing a 1mL aliquot. We performed population WGS by inoculating frozen populations into the evolution media and incubating for 24 hours, then removing an aliquot for DNA extraction for planktonic populations and sonicating a colonized bead and removing an aliquot for biofilm populations. We performed population WGS at day twelve for each lineage. We also isolated clones with colony morphologies and phenotypes distinct from the ancestor and sequenced these clones by WGS. Genotypes of evolved clones are shown in Table S1.

### Whole Genome Sequencing and Analysis

We extracted DNA using the DNeasy Blood and Tissue Kit (Qiagen, Hiden, Germany) and prepared the sequencing library as previously described (65, 66) using the Illumina Nextera kit (Illumina Inc., San Diego, CA). We sequenced populations to an average read depth of 86-254x and clones to an average read depth of 10-58x using an Illumina NextSeq500. Sequences were trimmed using the trimmomatic software v0.36 with the following criteria: LEADING:20 TRAILING:20 SLIDINGWINDOW:4:20 MINLEN:70 (67). The breseq software v0.35.0 was used to call variants using the −p polymorphism flag when analyzing population sequences (68). These parameters detect variants at >5% frequency within a population. To minimize false positive variant calls, we implemented breseq parameters requiring that variants be supported by at least three reads on each strand. For our reference genome, we used the *P. aeruginosa* UCBPP-PA14 109 genome (NC_008463) with annotations from the *Pseudomonas* Genome DB (69). Mutations that were detected between our ancestral strain and reference genome were removed from subsequent analysis as they were not selected by the experimental conditions. Variant calls indicative of poor read mapping were also removed, including variants that occurred at only the ends of reads, only within reads with many other mutations, or only at low total read depth. The breseq software also reports new junctions within the genome which may occur due to mobile element insertions, large deletions, structural rearrangements, and prophage excision and circularization, which we included in our analysis. Unfiltered breseq output and code documenting all analysis steps can be accessed at https://github.com/michellescribner/pa_nutrient. Filtering and plotting was performed in R v4.0.5 using packages tidyverse and ggrepel (70–72).

### Analysis of *morA* Mutations Detected in Laboratory and Clinical Environments

To identify instances of residue and domain-level parallelism, we analyzed nonsynonymous mutations in *morA* detected in this experiment, other laboratory experiments (23, 44), and clinical isolates (2). We determined the locations of *morA* mutations using the supplementary data of two studies (2, 23) and by analyzing sequences of populations propagated in the absence of antibiotic using breseq for Sanz-García et al. (44).

We also visualized *morA* mutations detected in other evolution experiments from our laboratory. These mutations were detected in an evolution experiment identical to this study but with the following alterations: populations were cultured in deep well plates with daily transfer through either 1:100 dilution or transfer of a colonized pipette tip into a well containing 5mL fresh media. Deep well plates were incubated with shaking at 100rpm. Three evolved lineages from each treatment were sequenced following twelve days of transfer. Whole population genome sequence data for these evolved populations are available in NCBI BioProject PRJNA692838.

The domain positions within the MorA protein were determined from the Pfam database (73). All nonsynonymous mutations detected in clinical and laboratory studies were visualized on the protein structure of PAO1 MorA published by Phippen et al. in Figure S1 (74, 75). Mutations were visualized based on their relative position within the PAO1 MorA protein using UCSF Chimera with color corresponding to domain (76).

### Colony Morphology

Colony morphologies of evolved mutants were visualized on agar plates containing 1% tryptone, 20ug/L Coomassie brilliant blue, and 40ug/mL Congo Red. Strains were grown overnight in the evolution media and 5uL was spotted onto plates. Plates were then incubated for 24 hours at 37°C followed by 72 hours at room temperature before photographing.

### Motility and Biofilm Assays

Motility and biofilm assays were performed similarly to as previously described with the alterations noted below (77–79). For swimming motility assays, evolved mutants were inoculated in the evolution media and incubated overnight. A sterile pipette tip was dipped into cultures and used to stab 0.3% agar supplemented with an M8 base, glucose, casamino acids, and MgSO4. Plates were incubated for 24 hours at 37 °C and diameter of the resulting growth was measured. Swarming motility assays were performed analogously to swimming assays, except plates contained 0.5% agar and 2.5uL of culture was spotted onto each plate. Plates were incubated for 24 hours at 37 °C then photographed. We estimated biofilm production of evolved mutants by crystal violet assay (79). We diluted overnight cultures of evolved mutants 1:100 in fresh minimal media to a volume of 200uL in a 96 well plate. After incubation for 24 hours at 37°C with shaking for 30 seconds every 15 minutes, we gently rinsed plates twice with water. We stained wells with 250uL of 0.1% crystal violet, incubated for 15 minutes, rinsed three times with water, then allowed them to dry overnight. Crystal violet was solubilized by adding 250 μl 95% EtOH solution (95% EtOH, 4.95% dH2O, 0.05% Triton X-100) to each well for 15 minutes. Biofilm formation was then visualized by measuring Abs OD_600_. Datapoints are the average of technical replicates from at least three independent experiments.

### Fitness assays

Evolved mutants were incubated for 24 hours in the evolution media, then inoculated into fresh tubes at the indicated starting proportions with a competitor (25uL of competitor A and 25uL of competitor B in 5mL fresh evolution media to generate a starting proportion of 0.5). Cultures were serially diluted in PBS and plated onto ½ concentration tryptic soy agar at the starting timepoint to estimate precise initial proportions. Cultures were then incubated for 24 hours, diluted 1:100 into fresh media, and incubated for another 24 hours. At the 48-hour timepoint, cultures were again plated. Fitness was calculated as selection rate as described by Turner et al (53) where A and B represent competitors A and B and d=0 and d=2 denote CFU/mL at timepoints 0 and 48 hours. Strains were differentiated by competing against a Lac+ marked ancestor that appears blue on agar containing X-Gal or differentiated by colony morphology in competitions of *morA* versus *lasR* mutants.

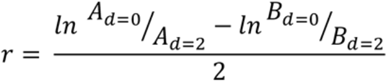

### Growth Curves

We measured growth curves of evolved mutants by growing cultures overnight in the evolution media, then diluting 1:100 into 200uL of fresh media in a 96 well plate. Cultures were incubated for 24 hours at 37 °C with shaking for 30 seconds every 15 minutes, and OD_600_ was measured at 15-minute intervals. We analyzed growth curves from at least three independent experiments with three replicates each. Growth curves were subsequently analyzed in R using the Growthcurver package (80).

### Nutrient Analysis

The ancestral strain and *lasR* R216Q mutant were grown from freezer stocks in the evolution media for 24 hours at 37°C. Cultures were diluted 1:100 into fresh media supplemented with all carbon sources (glucose, lactate, amino acids) and incubated at 37°C. We removed 1mL aliquots from these cultures every hour for 8 hours and collected supernatants by centrifugation at 13,000g for 1 minute, then filtered using a 0.2 mm filter. Two biological replicates for the ancestor and three for the lasR mutant were analyzed for nutrient content. We measured cell density by Abs OD_600_ for each aliquot prior to centrifugation.

Amino acid levels in the supernatants were analyzed using the Waters AccQ-Tag chemistry package. Samples were hydrolyzed using trichloroacetic acid precipitation. A 1:4 ratio of TCA was added to the samples, which were chilled on ice for 10 minutes and then centrifuged at 13,000 g for 10 minutes. The supernatant from the TCA precipitation was removed and pH balanced to pH 8.2-10 using KOH. The sample was derivatized using the AccQ-Tag methodology (Waters) and analyzed on a Waters HPLC system consisting of an e2985 Separations Module and a 2475 FLR Detector as per manufacturer’s instructions. For glucose and lactate consumption, samples were enzymatically analyzed using a D-Lactic Acid/L-Lactic Acid Enzymatic Bioanalysis UV-Test kit and a D-Glucose Enzymatic Bioanalysis UV-Test kit (Roche Diagnostics) with a modified manufacturer’s protocol to reduce total sample size to 300 uL. Results were read at 340 nm on a BioTek Synergy HTX plate reader.

### Statistical analysis

Data was analyzed using GraphPad Prism 9 and using R where noted.

### Data Availability

Data and code used for data analysis can be accessed at https://github.com/michellescribner/pa_nutrient. All sequencing reads were deposited in NCBI under BioProject accession numbers PRJNA595915 and PRJNA692838.

## Acknowledgments

We thank Catherine Armbruster and Michelle Clay for thoughtful feedback related to this work and the Microbial Genome Sequencing Center (MiGS) for genome sequencing. This work was supported by the National Institute of Allergy and Infectious Diseases at the National Institutes of Health (grant U01AI124302 and grant T32AI049820) and by the Cystic Fibrosis Foundation Research Development Program.

---

Figure S1. Mutated residues within MorA detected in this experiment, other laboratory experiments (23, 44), and clinical isolates (2) are illustrated on the structure of the PAO1 MorA DGC and PDE domains (PDB 4RNF) (74).

Figure S2. Mutations in the *lasR*, *morA*, and *wspF* genes produce rapid morphological diversity over short timescales. Images show representative morphologies of evolved mutants following incubation on agar supplemented with Coomassie Blue and Congo Red dyes. Mutations listed are the only variants detected between the isolated clone and the ancestor. Certain mutations identified in clones were not detected by population sequencing, thus likely represent rare haplotypes within the evolved lineages.

Figure S3. Decreased swarming motility of all clones isolated from evolved populations. Isolated clones from A) planktonic populations and B) biofilm populations were used to inoculate 0.5% agar plates. Plates were photographed after 24 hours of incubation.

Figure S4. Growth curves of evolved clones in different carbon sources reveal distinct growth advantages for mutants with *morA* and *lasR* alterations. A) Growth curves were performed in the media used for the evolution experiment (All Carbon) as well as media containing only a subset of the carbon sources (Glucose Only, Amino Acids Only, Lactate Only). Datapoints represent the mean of three independent experiments and error bars represent 95% CI. Three distinct *lasR* mutants and *morA* mutants were utilized for each growth curve assay. B) Area under curve, carrying capacity (k), and maximum growth rate (r) were determined for the growth curves visualized in A. Mean and 95% CI for the ancestor (gray), *lasR* mutants (blue), and *morA* mutants (green) are shown. Data were analyzed by two-way ANOVA with Tukey’s multiple comparisons, groups labeled with the same letter are not statistically different.

Figure S5. *LasR* mutations increase growth yield in independent culture. Clones isolated from planktonic cultures were incubated in the conditions of the evolution experiment for 24 hours then plated for CFU. Data from two independent experiments are shown with mean and 95% CI. Data were analyzed by one-way ANOVA (F(6,35) = 10.74, p < 0.0001) with post hoc comparisons between mutants and the ancestor using Dunnett’s multiple comparisons test (p < 0.05).

Figure S6. HPLC analysis of supernatants from a *lasR* R216Q mutant and the ancestral strain reveal that nutrients are consumed at similar rates between the two genotypes. Despite a spike in amino acid level at early timepoints for the *lasR* mutant, amino acids are consumed at similar rates as in the ancestral strain. Data represents the consumption of nutrients from media containing all carbon sources (amino acids, glucose, and lactate). Dynamics of amino acid levels for A) all amino acids and B) a representative amino acid (Aspartic acid) are shown. Consumption of C) glucose and D) lactate also occurs at similar rates between genotypes. Mean and SD are shown for at least two biological replicates. The slopes of nutrient consumption for the ancestral strain and *lasR* mutant were compared by t-tests controlled for false discovery rate (Benjamini-Hochberg, q = 5%), and no statistically significant differences were found.

Figure S7. The negative frequency dependence relationship between *lasR* and *morA* mutants persists across similar genotypes. We performed direct competitions at the starting concentrations indicated and measured relative fitness at 48 hours for a large deletion mutant including *lasR* and a *morA* mutant. Genotypes were differentiated by colony morphology. Data was analyzed by linear regression as shown, shaded area represents 95% CI of the regression. Slope = −0.9433, slope was statistically different from zero, F = 66.50, DFn = 1, DFd = 21, P<0.0001.

**Table S1.** Strains used in this study are shown with BioProject and BioSample numbers as well as mutations identified by our variant calling pipeline.

| Bioproject | Accession | Sample Name | Organism | Population of Origin | Variants |
| --- | --- | --- | --- | --- | --- |
|  |  | Ancestral Strain | PA14 |  |  |
| PRJNA692838 | SAMN17372736 | MRS 1301 | PA14 | Planktonic,<br>Population 1 | *lasR* R216Q |
| PRJNA692838 | SAMN17372737 | MRS 1302 | PA14 | Planktonic,<br>Population 2 | *morA* N1124K |
| PRJNA692838 | SAMN17372738 | MRS 1303 | PA14 | Planktonic,<br>Population 3 | *morA* K1123E |
| PRJNA692838 | SAMN17372739 | MRS 1304 | PA14 | Planktonic,<br>Population 4 | *morA* L1155Q |
| PRJNA692838 | SAMN17372740 | MRS 1305 | PA14 | Planktonic,<br>Population 4 | *lasR* V132G *morA*<br>L1155Q |
| PRJNA692838 | SAMN17372741 | MRS 1306 | PA14 | Planktonic,<br>Population 5 | *lasR* R216Q *pf5r*<br>16bp DUP |
| PRJNA692838 | SAMN17372742 | MRS 1307 | PA14 | Planktonic,<br>Population 5 | *morA* L1155Q<br>*pf5r* 16bp DUP |
| PRJNA692838 | SAMN17372743 | MRS 1308 | PA14 | Biofilm,<br>Population 2 | PA14_64050<br>R26C *lasR* P117L |
| PRJNA692838 | SAMN17372744 | MRS 1309 | PA14 | Biofilm,<br>Population 2 | *wspF* Δ1bp |
| PRJNA692838 | SAMN17372745 | MRS 1310 | PA14 | Biofilm,<br>Population 3 | *wspF* L330Q |
| PRJNA692838 | SAMN17372746 | MRS 1202 | PA14 | Planktonic,<br>Population 2 | PA14_45920..PA1<br>4_46440 Δ49,133<br>bp |
| PRJNA692838 | SAMN17372747 | MRS 1203 | PA14 | Planktonic,<br>Population 3 | PA14_45800..PA1<br>4_46240 Δ40,707<br>bp |
| PRJNA692838 | SAMN17372748 | MRS 1219 | PA14 | Biofilm,<br>Population 1 | PA14_64050<br>R26C *pilC* Δ5bp |
| PRJNA692838 | SAMN17372749 | MRS 1220 | PA14 | Biofilm,<br>Population 1 | *lasR* G164* |
| PRJNA692838 | SAMN17372750 | MRS 1221 | PA14 | Biofilm,<br>Population 1 | *wspF* E189* *pilC*<br>Δ1bp |
| PRJNA595915 | SAMN14929132 | MRS_1021_1 | PA14 | Planktonic,<br>Population 1 | *morA* E1153K<br>*tRNA<sup>Thr</sup>/tufB*<br>(+20/65) G->A |

**Supplementary Data 1.**

Sheet 1: Average read depth for evolved populations.

Sheet 2: SNPs and small indels in evolved populations.

Sheet 3: New Junction Evidence in evolved populations.

**Supplementary Data 2.**

Sheet 1: Average read depth for evolved clones.

Sheet 2: SNPs and small indels in evolved clones.

Sheet 3: New Junction Evidence in evolved clones.

